# Fast and accurate defocus for improved tunability of cryo-EM experiments

**DOI:** 10.1101/2019.12.12.874925

**Authors:** Radostin Danev, Hirofumi Iijima, Mizuki Matsuzaki, Sohei Motoki

## Abstract

Current data collection strategies in electron cryo-microscopy (cryo-EM) record multiframe movies with static optical settings. This limits the number of adjustable parameters that can be used to optimize the experiment. Here, we propose a method for fast and accurate defocus (FADE) modulation during movie acquisition. It uses the objective lens aperture as an electrostatic pole that locally modifies the electron beam potential. The beam potential variation is converted to defocus modulation by the typically undesired chromatic aberration of the objective lens. The simplicity, electrostatic principle and low electrical impedance of the device will allow fast switching speeds that will enable per-frame defocus values in cryo-EM movies. Researchers will be able to define custom defocus “recipes” and tailor the experiment for optimal information extraction from their sample. The FADE method could help to convert the microscope into a more dynamic and flexible optical platform that delivers better performance in cryo-EM single particle analysis and electron cryo-tomography.

## 1. Introduction

The introduction of direct electron detectors (DEDs) revolutionized the cryo-EM field. With their improved signal-to-noise ratio and multiframe movie capabilities, these new cameras led to the “resolution revolution” that propelled cryo-EM from a niche technique to a powerful mainstream structural method (Rivera-Calzada & Carroni, 2019). Cryo-EM methodology continues to improve, with advancements in many areas, such as image analysis (Zivanov *et al.*, 2019*b*), data acquisition strategies (Hagen *et al.*, 2017; Cheng *et al.*, 2018; Chreifi *et al.*, 2019; Eisenstein *et al.*, 2019), on-the-fly image processing (Tegunov & Cramer, 2019), detectors (Song *et al.*, 2019; Mendez *et al.*, 2019; Feathers *et al.*, 2019), sample preparation (Dandey *et al.*, 2018; Ravelli *et al.*, 2019; Rubinstein *et al.*, 2019; Kontziampasis *et al.*, 2019) and automation (Schorb *et al.*, 2019). On the optics side, recent developments include the introduction of cold field emission guns (Hamaguchi *et al.*, 2019), the use of lower accelerating voltages (Wu *et al.*, 2019; Naydenova *et al.*, 2019) and the development of a laser phase plate (Schwartz *et al.*, 2019).

The movie acquisition capability of DEDs enables alignment and registration of individual movie frames to counteract one of the main resolution-limiting factors in cryo-EM, the beam-induced specimen motion. When irradiated, frozen-hydrated cryo-specimens experience changes in their shape that cause image blurring (Brilot *et al.*, 2012), presumably due to release of mechanical stresses built up during the freezing process (Thorne, 2019). Alignment and averaging of the movie frames, called motion correction, can minimize the blurring and its dampening effect on high-resolution components (Zheng *et al.*, 2017).

Another important use of the fractionated exposures from DEDs is maximizing the signal-to-noise ratio of images by considering the effects of radiation damage on the sample. Averaging methods, such as per-frame exposure weighting (Grant & Grigorieff, 2015) and Bayesian polishing (Zivanov *et al.*, 2019*a*), improve the spectral signal-to-noise by weighted filtering of individual frames that takes into account the fading of high-frequency components as the sample is irradiated. At present, motion correction and exposure weighting are the sole applications of the multi-frame capabilities of DEDs in cryo-EM. With this work, we hope to expand the utilization of the third dimension in the movies and improve the flexibility of experiments.

Phase contrast is essential for optimal information extraction in cryo-EM (Glaeser, 2019). Beam-sensitive low atomic weight biological samples modulate predominantly the phase and much less the amplitude of the electron wave. Out-of-focus image acquisition, i.e. with defocus, is the main phase contrast mechanism in cryo-EM. There have been other proposed phase contrast methods, such as phase plates (Danev & Baumeister, 2017), but to date the defocus method has produced the highest resolutions and remains the most convenient approach (Danev *et al.*, 2019). Despite optimistic theoretical estimates, and similarly promising experimental tests (Russo & Henderson, 2018), in practice, the defocus approach dictates a compromise between contrast and resolving power. Higher defocus values produce stronger contrast but limit the achievable resolution. Low defocus values extend the high frequency information coverage, but the images have very weak contrast. In general terms, the defocus setting acts as a tunable bandpass filter which can be used to emphasize either high or low periodicity features of the sample. Single particle experiments use a range of defocus values to get better spectral coverage. For many years, this has been the de facto standard data acquisition method in cryo-EM and it continues to produce outstanding results.

The present cryo-EM imaging technique uses static optical parameters to collect multiframe movies. The defocus value, astigmatism setting, and beam tilt angle are kept constant throughout the exposure. The only variables are the beam-induced motions of the specimen and the progressive radiation damage of the sample, which causes gradual fading of high-resolution components. Theoretically, the overall signal-to-noise ratio of the micrograph could be maximized by an optical filter that follows the information content of the sample through the exposure. Defocus is the obvious candidate for such a filter because of its bandpass characteristics.

In this work, we investigate a novel approach for fast and accurate defocus (FADE) modulation with the goal of achieving real-time control of the optical properties of the microscope. Such capability will improve the tunability of the experiment and provide additional means for maximizing the performance of the two main cryo-EM methods, single particle analysis and electron cryo-tomography. High-speed modulation, preferably in sync with the hardware framerate of the detector, is needed to minimize the defocus transition time, which will be lost in terms of image information. Current generation DEDs operate at more than one kilohertz framerate and therefore transition times shorter than one millisecond are desirable. The accuracy of the modulation is also very important. Due to sample geometry variations, the base defocus of a micrograph is unknown and must be determined during image processing through a process called contrast transfer function (CTF) fitting (Rohou & Grigorieff, 2015). Adding a modulation pattern on top of the base defocus should not introduce additional unknown variables that would reduce the accuracy of the CTF fits. The defocus offsets must have high absolute certainty, in the order of a nanometer, so that the only unknown variable remains the base defocus. In practice, the modulation patterns, referred here as “modulation recipes”, will consist of defocus versus time (or detector frame number) series that could be freely adjusted by the practitioner to best suit the needs of the experiment.

There are several practical ways to control the defocus in real time, some of which have already been investigated. The most obvious and direct approach is to use the objective lens of the microscope. This approach has two main limitations, both related to the fact that the objective is a strong magnetic lens. The high electrical impedance of the lens coil precludes fast modulation and the magnetic hysteresis of the lens polepiece, which operates near magnetic saturation, will affect the accuracy of the modulation. Another practical option for defocus modulation is through variation of the accelerating voltage of the microscope, which effectively changes the wavelength of the electrons. This approach has been proposed and successfully tested in practice for high-resolution real-time imaging of materials science samples (Ando *et al.*, 1994; Takai *et al.*, 2005). It satisfies both criteria for high-speed and high-accuracy modulation, however, because the modulation is applied on the beam energy it influences the behavior of the whole optical system. In relation to cryo-EM, this has a practical implication about the usability of energy filtering in such a system. Energy filters are widely used in cryo-EM because they improve the signal-to-noise-ratio by removing inelastically scattered electrons from the image (Yonekura *et al.*, 2006; Sigworth, 2016). In such applications, the filter is operated in zero-loss energy filtering mode with a narrow energy selection slit. If the accelerating voltage is rapidly modulated, the energy filter must follow the primary beam energy in order to maintain the zero-loss filtering mode. This will be quite challenging in practice due to the optical characteristics and the construction of the filter (Reimer, 1995).

Here, we propose a defocus modulation method based on an additional optical element at the back-focal plane of the objective lens that is independent from the rest of the optical system of the microscope. The back-focal plane, also called diffraction plane, is located at the bottom of the objective lens polepeiece gap and is the place where objective lens apertures or phase plates are typically applied. Through optical simulations, we investigated several modulator designs and tested in practice the simplest solution based on electrostatic biasing of the objective lens aperture. The miniature size and electrostatic nature of such modulator help to satisfy the high speed and high accuracy FADE requirements. Its simple construction will also make it an easy and cost-effective upgrade for existing microscopes.

## 2. Materials and methods

### 2.1 Optical simulations

The simulation of the electrostatic potential distribution and focusing power of an electrostatic micro-lens was performed in Wolfram Mathematica software (Wolfram Research, IL, USA). The potential distribution was determined by numerically solving the cylindrical coordinate form of Laplace’s equation on a mesh. The focal length of the lens was estimated by numerically solving the paraxial ray equations for rotationally symmetric optical elements (eqs. (15.38-15.40) in (Hawkes & Kasper, 2017)). To validate the numerical results, we compared them to approximate expressions for Einzel lenses (eqs. (11) and (31) in (Hawkes *et al.*, 1995)). The values were within 10% of each other.

The parameters of the magnetic micro-lens design were estimated using a bell-shaped field thin-lens approximation formula (eq. (2.9) in (Egerton, 2016)) and numerical integration of the Bush’s formula for a thin weak-field lens (eq. (17.78) in (Hawkes & Kasper, 2017)). The results were within 2% of each other.

The numerical simulation of the electrostatic potential distribution around a voltage-biased objective lens aperture was performed with ElecNet v6 software (Mentor Graphics, OR, USA). The magnetic field distribution inside the objective lens polepiece gap was numerically calculated with MagNet v6 software (Mentor Graphics, OR, USA). Calculations of electron trajectories inside the objective lens polepiece gap with a biased aperture were performed by numerically solving the paraxial ray equation (eq. (15.10) in (Hawkes & Kasper, 2017)) with custom Python scripts. The optical parameters and objective lens area dimensions for the simulations were taken from the microscope used for the experiments described below.

### 2.2 Experimental measurement of defocus modulation

Experimental measurement of the effect of objective lens aperture bias on the defocus were performed on a JEOL JEM-2200FS (JEOL, Akishima, Japan) electron microscope operated at 200 kV accelerating voltage. It was equipped with an objective lens cryo-polepiece (CRP) with *f*_0_ = 2.8 mm focal length, *C*_s_ = 2.0 mm spherical aberration and *C*_c_ = 2.1 mm chromatic aberration coefficients. An Au/Pd diffraction grating replica test sample was used to produce strong power spectrum signals for more accurate contrast transfer function fitting. The objective lens aperture holder’s grounding wire was disconnected from ground and was connected to a variable DC power supply through a 10 kOhm protection resistor. The bias voltage on the aperture was varied in 10 V steps between 0 and 100 V, except for image series 1, where the maximum applied voltage was 60 V. The holder was inserted in the beam path at a 150 μm objective aperture position. Images were acquired at x200K indicated magnification and calibrated pixel size of 0.44 Å/pix with a Gatan US1000XP (Gatan, CA, USA) CCD camera. The defocus and astigmatism parameters of the voltage series images were estimated with CTFFIND4 software (Rohou & Grigorieff, 2015).

## 3. Results

### 3.1 Overview of the FADE system

A principle schematic of the proposed FADE microscope system is shown in Fig.1. The defocus modulator is located below the sample and the objective lens. It is connected to the FADE control unit, which generates a synchronized defocus modulation pattern and applies it during an exposure. The FADE control unit contains a digital modulation controller which receives synchronization input from the direct electron detector in the form of beam shutter signal or frame timing pulses. The controller outputs the pattern as a digital stream to a high-precision digital-to-analog converter (DAC). The analog output of the DAC is connected to a high-voltage linear amplifier that amplifies the signal to the appropriate amplitude for driving the modulator. The defocus recipe is uploaded to the FADE controller by the data acquisition control computer. The recipe consists of a sequence of defocus values versus time or DED frame number.

**Figure 1.**
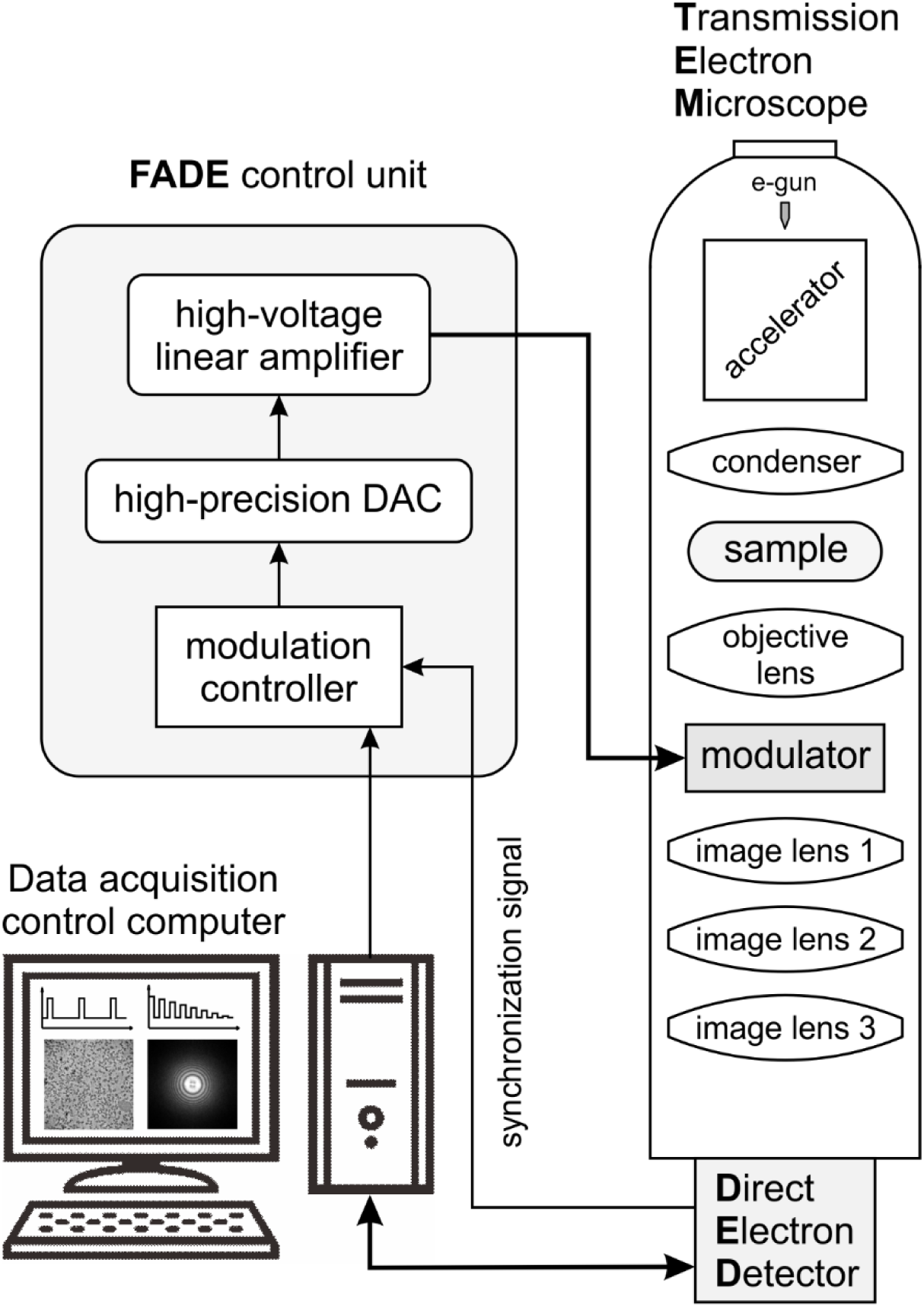
Overview of the fast and accurate defocus (FADE) microscope system. The FADE control unit receives synchronization signal from the direct electron detector and applies a pre-programmed defocus modulation pattern to the modulator. Defocus modulation recipes are uploaded to the FADE unit from the data acquisition control computer.

### 3.2 Micro-lens optical setup

Physically, the FADE modulator could be a micro-lens positioned at the back-focal plane of the objective lens. Fig. 2 illustrates such optical arrangement. The sample is located very close to the front focal plane of the objective lens, which has a focal length *f*_0_. A simple geometrical optics calculation shows that to produce defocus Δ*f*, the focal length of the micro-lens *f*_m_ must be *f*_m_=*f*_0_^2^/Δ*f*. For example, in a typical cryo-microscope with objective focal length of 3.5 mm the micro lens must have a focal length of ~12 m to produce a defocus of 1 μm. This shows that the micro-lens is a weak lens, but as demonstrated by the simulations below, even a weak lens would be challenging to produce in the miniature form required to fit in the available space at the back-focal plane. The focal length of the micro-lens is proportional to the square of the objective focal length. For strong narrow-gap objective lenses, like those typically used in materials science, the micro lens would have to be several times more powerful to produce the same amount of defocus.

**Figure 2.**
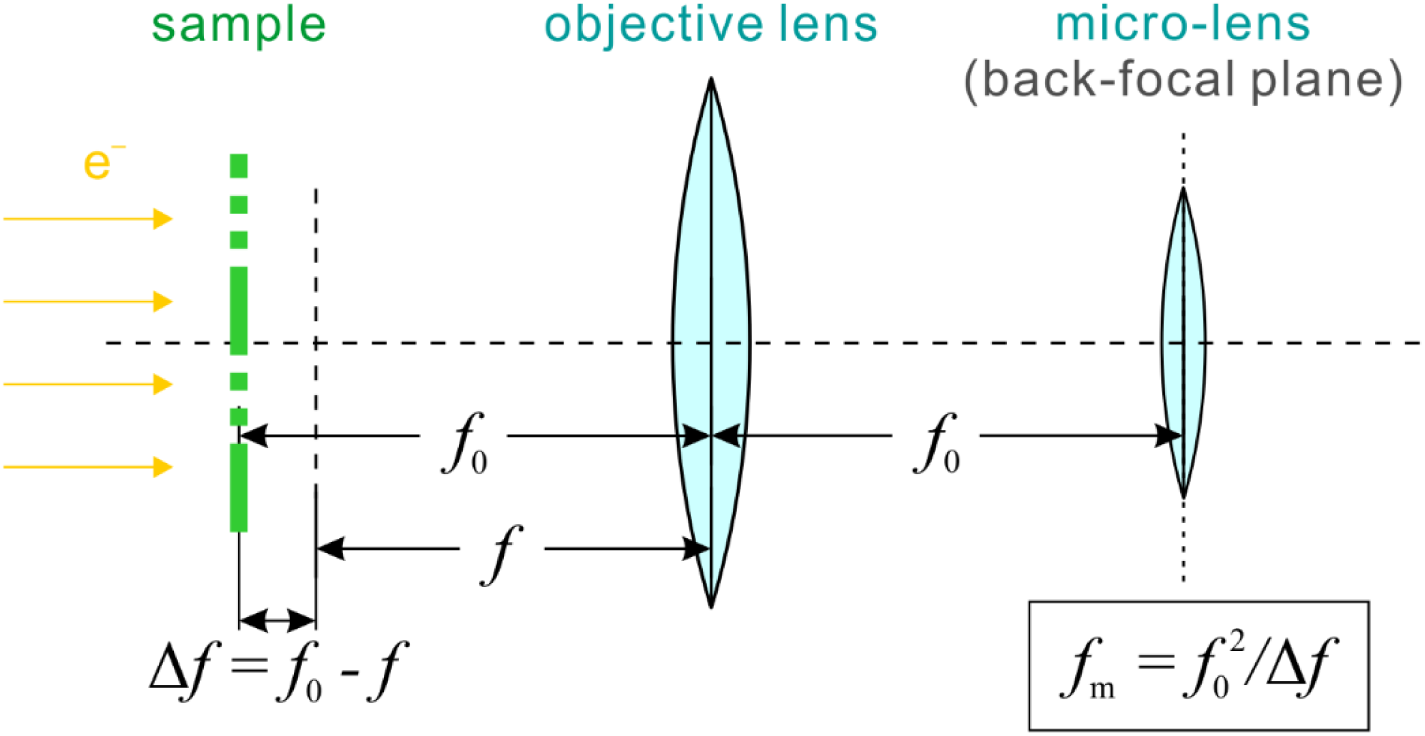
Optical diagram of the sample, the microscope objective lens and a modulator mirco-lens arrangement. The micro-lens is positioned at the back-focal plane of the objective lens; the same location where objective lens aperture is typically used. The formula below the mico-lens describes the geometrical optics relation between the sample defocus Δ*f*, the objective lens focal length *f*_0_ and the micro-lens focal length *f*_m_.

### 3.3 Electrostatic micro-lens (Einzel lens) design

We simulated numerically the performance of an electrostatic micro-lens (Fig. 3a) to determine the required excitation voltages for achieving a desired lens power. Fig. 3c shows the electrostatic field distribution inside a three electrode Einzel lens. The outer electrodes are kept at ground potential and act as shields. The central electrode is connected to the control voltage. Using the above ~12 m estimate for the required focal length of the micro-lens and 200 kV primary electron energy, the Einzel lens simulation showed that the central electrode must be set to a potential of ~1600 V. Considering the small dimensions of the lens and the narrow separation between the electrodes (~0.2 mm), such a high potential may be difficult to achieve in practice. It would require dielectric strength of at least 10 MV/m between the electrodes.

**Figure 3.**
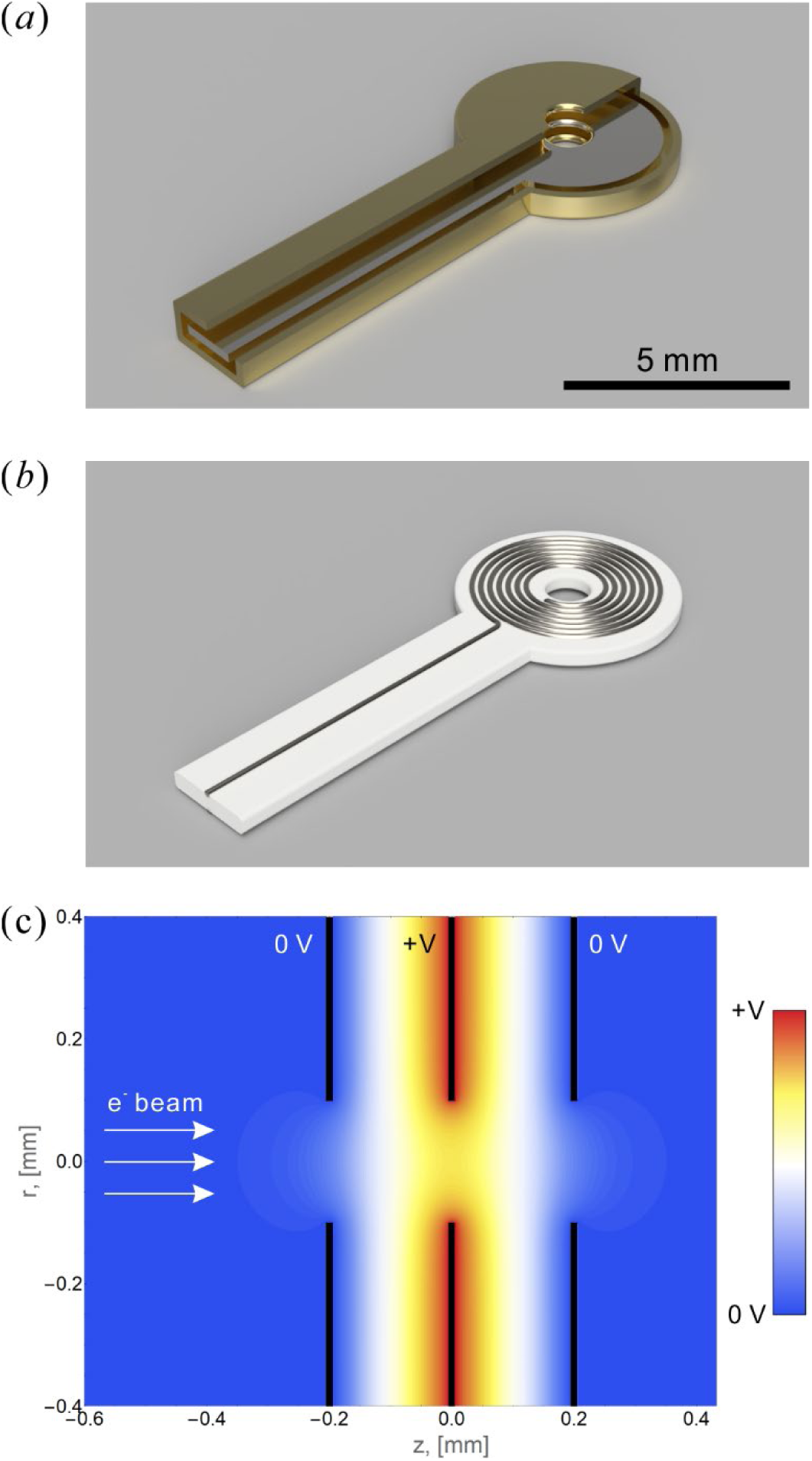
Conceptual 3D renderings of (a) electrostatic and (b) magnetic micro-lens designs. The electrostatic lens concept in (a) has a cutout showing the inner control electrode. The magnetic micro-lens in (b) consists of a flat metal coil on top of a non-conductive support. (c) Electrostatic potential distribution calculated through numerical finite element analysis of a three electrode (“Einzel”) electrostatic lens. The outer electrodes are kept at ground potential and act as shields. The control voltage is applied to the inner electrode. The electron beam passes through the inline circular openings in the electrodes.

### 3.4 Magnetic micro-lens design

Another possible design for the micro-lens is a flat magnetic coil (Fig. 3b). Calculations based on a thin and weak lens approximations showed that the micro coil must be energized with ~50 At (ampere-turns). In other words, if the coil has 50 turns the current must be ~1 A. Again, considering the small physical dimensions of the device, such current densities would be quite challenging to achieve in practice.

### 3.5 Objective lens aperture bias design

A third approach for defocus modulation is to apply bias voltage to the objective lens aperture. Fig. 4 contains an illustration of the component arrangement inside the objective lens polepiece gap. The sample holder is located inside a cryo-box that shields it and prevents the accumulation of water ice contamination on its surface. The objective lens aperture is located inside a ~1 mm gap between the cryo-box and the lower objective lens polepiece. During normal operation of the microscope, the objective lens aperture is grounded to prevent electromagnetic disturbances to the beam. For the FADE application, the aperture will be connected to a voltage generator and will act as a defocus modulator.

**Figure 4.**
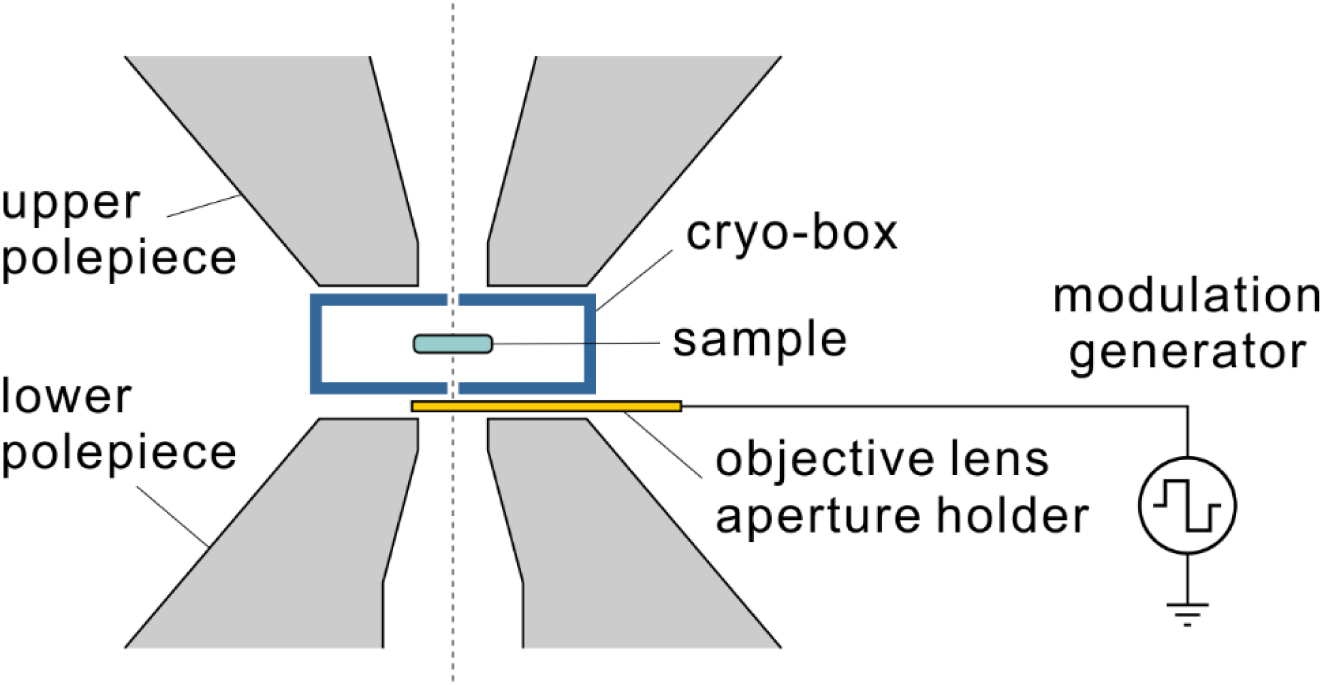
Sketch of the objective lens area of an electron cryo-microscope. The sample is located inside a cryo-box that minimizes water ice contamination buildup on its surface. The objective lens aperture holder is positioned between the cryo-box and the lower polepiece of the objective lens. By connecting the objective lens aperture to a voltage source, it can be used as a FADE modulator.

Magnetic lenses have chromatic aberration that expresses the fact that the lens has lower focusing power for higher energy electrons. The defocus Δ*f* induced by electron beam potential change Δ*U* is given by the formula (from eq. (66.38) in (Hawkes & Kasper, 1996)):

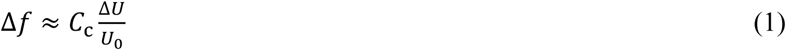

where *C*_c_ is the chromatic aberration coefficient of the lens and *U*_0_ is the primary accelerating voltage. The chromatic aberration coefficient of electron microscope objective lenses is in the order of a few mm and is similar in magnitude to the spherical aberration coefficient *C*_s_ and the focal length. The energy spread of the electron source combined with the chromatic aberration of the objective lens causes defocus spread, which is the dominating optical performance limiting factor in modern electron microscopes (Naydenova *et al.*, 2019). In this application case, however, we can exploit the typically undesirable chromatic aberration for constructive purposes. If an electric potential is applied to the objective aperture (Fig. 5a), the electron beam passing through the aperture will experience a change in potential equal to the applied voltage. The potential change will happen locally, in a region around the aperture holder and will diminish quickly away from the holder. The electron beam energy throughout the rest of the optical system will not change and therefore the operation of other optical elements, such as an energy filter, will not be affected. The local modulation of the electron beam potential combined with the chromatic aberration of the objective lens will produce a defocus change proportional to the applied voltage. Because the potential is applied only in a portion of the optical path through the lens, the defocus change will be smaller than that given by formula (1).

To more accurately estimate the defocus modulation with a biased aperture, we performed numerical simulations of the potential distribution around the aperture holder (Fig. 5a) and combined the results with the axial magnetic field profile of the objective lens (Fig. 5b). The numerical results showed a linear modulation profile with modulation strength of 5.87 nm/V (Fig. 5c). For the same optical setup (*C*c = 2.1 mm, *U*_0_ = 200 kV), formula (1) gives a modulation strength of 10.5 nm/V, which is approximately twice larger than the simulation result. This is in line with the fact that the modulation potential is applied mostly in the region between the bottom of the cryo-box and the lower polepiece (Fig. 5a).

**Figure 5.**
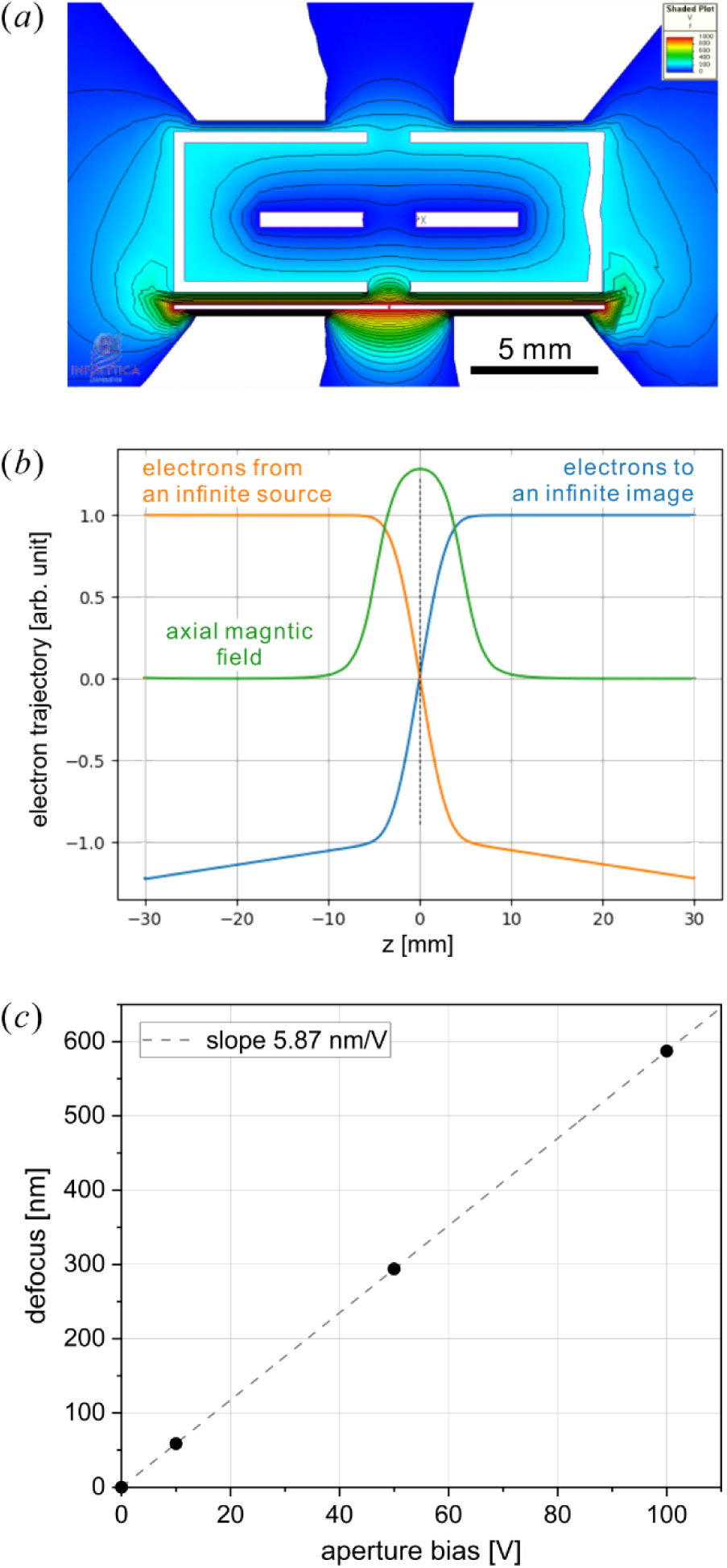
Results from a numerical simulation of the effect from a voltage bias on the objective lens aperture. (a) Electrostatic potential distribution around the objective lens polepiece gap caused by applying voltage to the objective lens aperture. The potential is confined mostly between the bottom of the cryo-box and the lower polepiece. (b) Axial magnetic field distribution along the optical axis (green) and electron trajectories for a paraxial illumination ray (orange) and an image ray (blue). (c) Numerical results for the defocus change caused by biasing the objective lens aperture. The results indicate a linear relation between defocus and bias voltage with modulation strength of 5.87 nm/V.

Results from experimental measurements of the defocus modulation at several aperture bias voltages are presented in Fig. 6. The defocus and astigmatism, calculated as the difference between the high and low defocus axes (Fig. 6b), were measured through CTF fitting of the power spectra (Fig. 6a) from three image series. The defocus and astigmatism modulations were calculated by subtracting the 0 V intercepts of the linear fits of the experimental data (Fig. 6c,d), which correspond to the base defocus and astigmatism of the objective lens. The defocus modulation exhibited a linear dependence on the applied voltage (Fig. 6e) with a slope of 4.29 ± 0.04 nm/V. The astigmatism modulation increased linearly with the applied voltage (Fig. 6f) with a slope of 0.953 ± 0.008 nm/V. There was a very good agreement and consistency between the measurements from the three image series (Fig.6e,f). We attribute the residual deviations of the experimental points from the linear curves to errors in the CTF fits, although this assertion could not be confirmed based on the current data.

**Figure 6.**
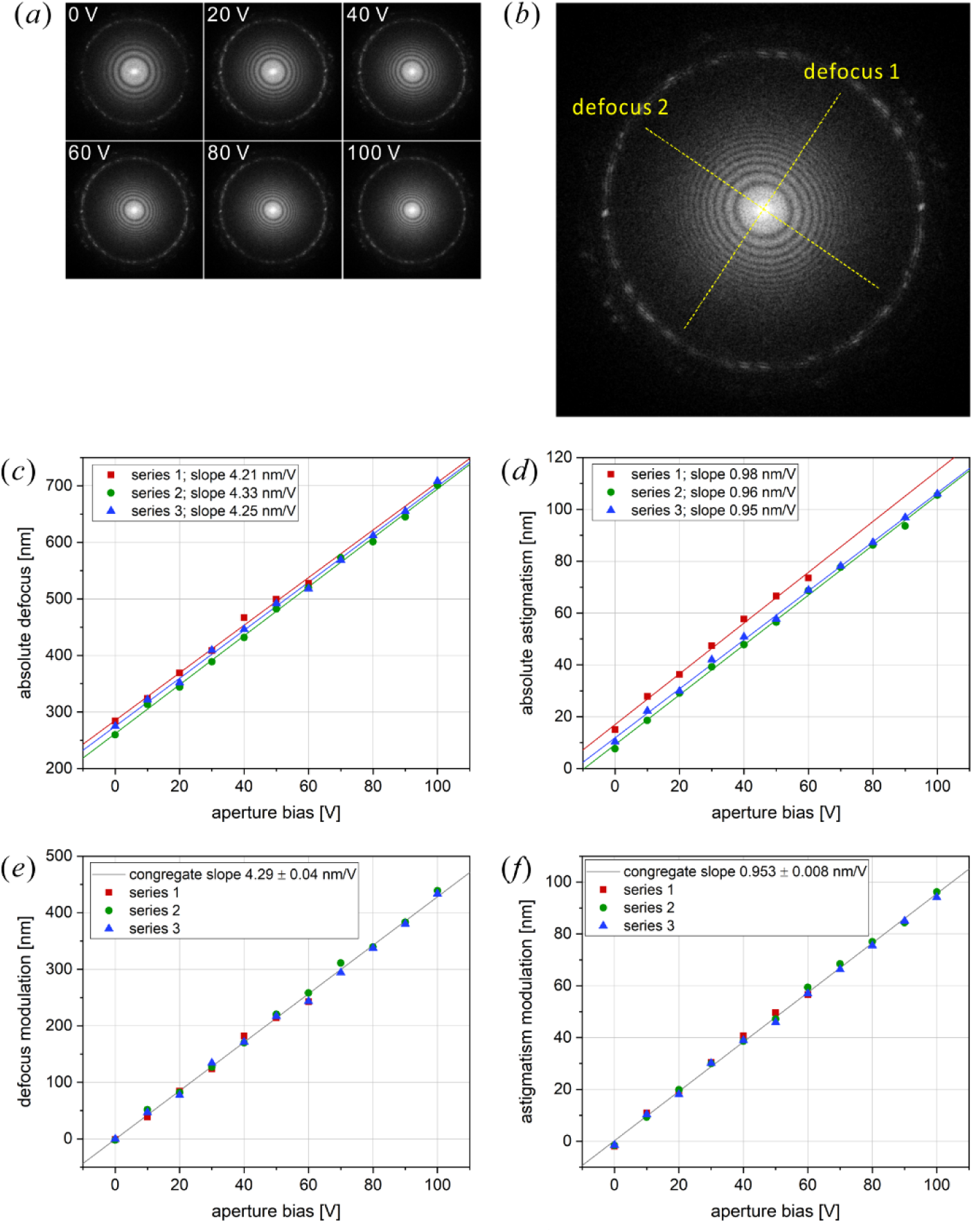
Experimental results from a FADE modulation test. (a) Power spectra of a series of images from a test sample collected at several aperture bias voltages. The contrast transfer function (CTF) Thon rings are moving inwards, indicating a defocus increase as the aperture voltage is increased. (b) The defocus values were measured by fitting the two-dimensional CTF profile in the power spectra. Different defocus values in perpendicular directions indicate the presence of image astigmatism, which is calculated as the difference between the maximum and minimum defocus axes, i.e. (defocus1 - defocus2). (c) Plot of the measured absolute defocus values as a function of modulation voltage. The results are from three independent image series. The lines are linear fits of the data with slopes shown in the legend. (d) Absolute astigmatism values with linear fits. (e) Defocus modulation calculated by subtracting the 0 V intercepts of the linear fits in (c) from the data points. The linear fit in (e) is for the congregation of data points with slope shown in the legend. (f) Astigmatism modulation calculated by subtracting the 0 V linear fit intercepts in (d) from the data points. The line is a congregate linear fit with slope indicated in the legend.

## 4. Discussion

The FADE approach proposed here will allow researchers to optimize their cryo-EM experiments by giving them the freedom to select an appropriate defocus modulation strategy. A few examples of modulation recipes are shown in Fig. 7. The simplest case is a defocus step after a predefined initial exposure to the sample (Fig. 7a). Due to radiation damage, high-resolution information fades quickly after the first few electrons per square angstrom of exposure. For example, the first ~10 e^−^/Å^2^ of exposure could have a relatively low defocus (e.g. ~200 nm) to maximize high resolution information capture followed by higher defocus (e.g. ~1000 nm) for the rest of the exposure to increase the overall contrast. The defocus step recipe will be the easiest to realize experimentally because it does not require continuously variable modulation voltage, just an on/off switch. It will also be the simplest to process by separating each movie into low and high defocus frame groups for CTF fitting and image processing.

**Figure 7.**
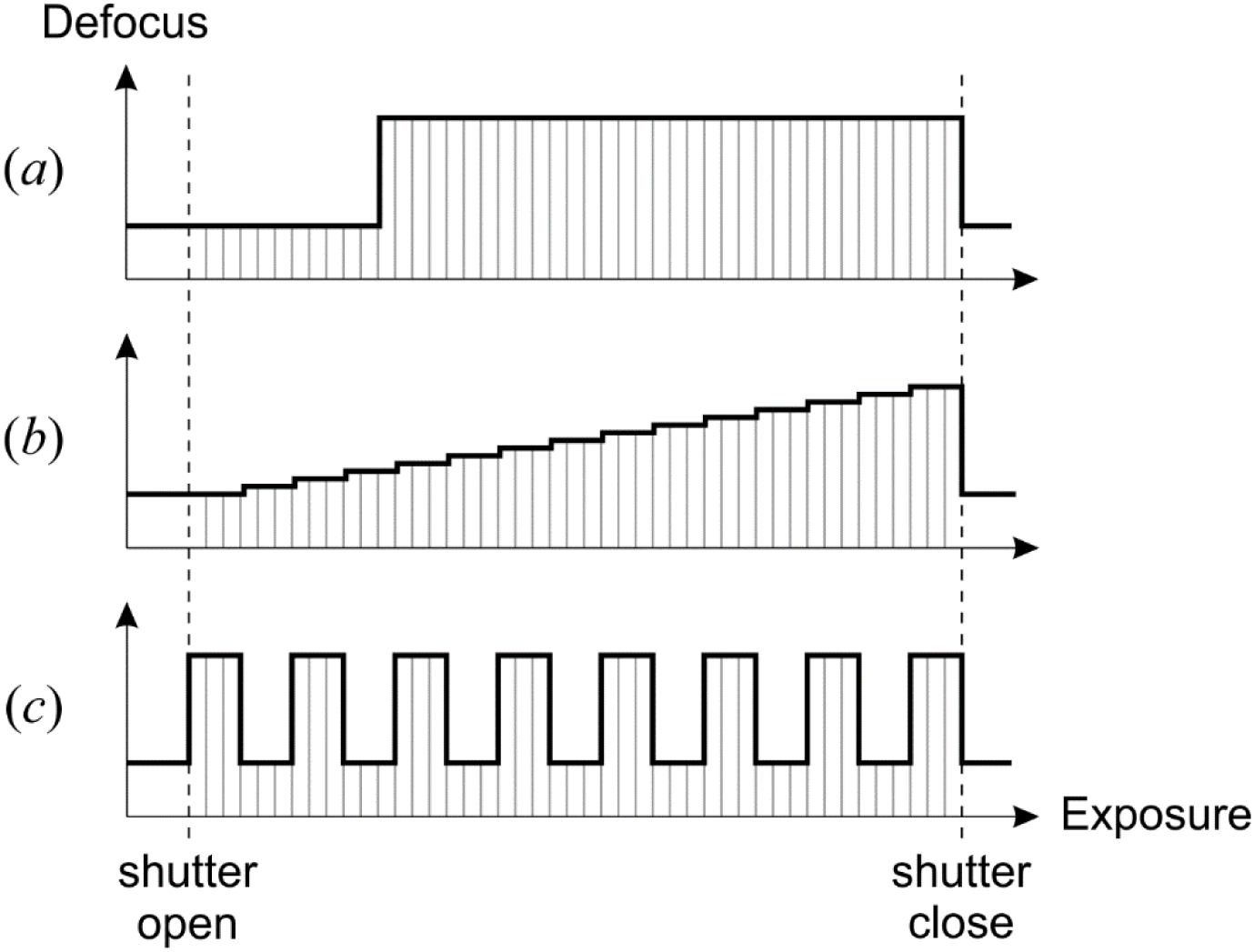
Defocus modulation recipe examples. (a) Defocus step: The movie starts with low defocus to preserve high resolution information from the undamaged sample and after an initial exposure the defocus jumps to a higher value for the rest of the movie for improved contrast. This recipe will be easy to implement and process because there are only two defocus values and just one switch point. (b) Defocus ramp: The defocus gradually increases throughout the exposure matching the damage level of the sample. This recipe will be more difficult to realize experimentally because it requires proportional voltage control. The data processing will also be more complicated because of the multiple defocus frame groups in each movie that have to be handled independently during CTF fitting and image demodulation. (c) Defocus oscillation: The defocus alternates between two values during the exposure. This recipe will be more suited for electron cryo-tomography with the higher defocus value intended to produce contrast and the lower to preserve high resolution information. It will be similar in difficulty to the defocus step recipe because it uses only two defocus values.

A more advanced recipe is shown in Fig. 7b. It is a linear approximation of a tuned filter that tries to maximize the information extraction from the sample as the exposure and radiation damage progress. The exposure starts at a low defocus that gradually increases throughout the movie. Although such a technique would theoretically provide an optimal result, it will be more challenging experimentally and especially for image processing. The defocus modulation will require accurate voltage control. The CTF fitting would have to be performed in a three-dimensional fashion, instead of the standard 2D power spectrum fits, with the third dimension being the modulation defocus. Data processing will also be more complicated with each particle having to be treated as a three-dimensional defocus stack instead of the two-dimensional images used nowadays.

The oscillating defocus recipe in Fig. 7c is more suited for electron cryo-tomography experiments. In cryo-tomography, the data consists of a series of movies acquired at different sample tilt angles. To keep the total exposure within reasonable limits, each tilt image is acquired with just 2-3 e^−^/Å^2^ distributed over ~10 frames. The relatively noisy tilt images must be aligned accurately to produce a high-resolution 3D tomogram. For this reason, and to increase the overall contrast in the tomogram, cryo-tomography requires higher defocus values (>2 μm) than single particle analysis (0.5-1.5 μm). The oscillating defocus scheme will provide high defocus movie frames (e.g. 3 μm) for increased contrast and accurate alignment and low defocus frames (e.g. 1 μm) for better preservation of the high-resolution information. The combination of the two defocus subsets will result a more uniform spectral coverage, which is usually lacking in cryo-tomography datasets. The oscillating defocus scheme will be relatively easy to implement because, as with the defocus step recipe, it only requires an on/off switch type voltage control. From processing viewpoint, the CTF fits and image handling will necessitate only two frame groups, one with low and one with high defocus, and therefore will also be similar in difficulty to the defocus step recipe.

The defocus recipes discussed above are just examples. The FADE method will provide infinite tunability and complete freedom in selecting the defocus profile. However, as mentioned for the staircase recipe (Fig. 7b), more complicated defocus profiles will require more sophisticated image processing strategies. Even for the simple step and oscillating profiles (Fig. 7a,c), the current image processing routines will have to be extended to support multiple defocus values during CTF fitting and for each particle/tilt image during demodulation. By design, one of the main advantages of the FADE method is the absolute accuracy of defocus modulation. The modulation offsets will be precisely known and therefore the number of variables that must be determined by CTF fitting (e.g. defocus and astigmatism) will remain the same as with the conventional constant defocus method. Nonetheless, depending on the mechanical and electrical stability of the hardware, the absolute modulation coefficients for defocus and astigmatism may have to be measured before each experimental session by collecting a few calibration movies with a thin carbon film specimen. Because of the smoothness of the local electrostatic potential distribution around the objective lens aperture, we don’t expect FADE modulation to have adverse effects on higher order aberrations of the microscope, but this remains to be confirmed experimentally.

The a priori known absolute defocus offsets could offer another experimental advantage – the use of very low defocus values that are currently impractical due to low contrast and insufficient CTF rings for accurate fits. For example, using the defocus step recipe in Fig. 7a the low defocus region could be collected with very low defocus of ~100 nm and the rest of the frames with ~1000 nm defocus. CTF fits and particle picking will be based on the high defocus portion of the movie. The low defocus part will only serve as a high-resolution information reservoir. It remains to be determined experimentally if beam-induced motion tracking will be possible with just the high-resolution information in the initial part of a movie. One possibility to mitigate motion tracking issues would be to interlace a few high defocus frames in this part.

Hardware wise, we evaluated three possible designs for FADE modulation. Simulation results showed that the required voltages and currents for electrostatic and magnetic mirco-lenses are close to the limits of what would be technically possible in practice. Such devices will be challenging to design and costly to manufacture and maintain. The objective aperture voltage bias approach is much simpler, and we expect that with minor hardware modifications it can be implemented in most existing microscope systems. Our simulations and experimental results confirmed the feasibility of this method and demonstrated that a sufficiently strong defocus modulation could be achieved in practice. The aperture bias method exploits the otherwise undesired chromatic aberration of the objective lens. It must be emphasized that the modulation effect is almost exclusively due to the chromatic aberration and not due to electrostatic lens formation between the aperture and the surrounding ground planes. The approach supports bipolar modulation, i.e. positive and negative bias voltage, that will double its modulation range. This can be used to overcome maximum voltage limits due to e.g. electrostatic isolation restrictions for applications requiring wider modulation ranges.

In terms of limitations and challenges, the aperture bias FADE is quite simple and robust. We measured experimentally a modulation strength of ~4.3 nm/V at 200 kV. Everything else being equal, on a 300 kV microscope the modulation strength will be (from eq. (1)) 300/200 = 1.5 times lower or ~2.9 nm/V. To achieve a desired defocus modulation amplitude of 1μm the applied aperture voltage must be ~230 V @ 200 kV and ~350 V @ 300 kV. These voltages are not in ranges that would be difficult to manipulate electronically and/or handle in practice. The stability and accuracy requirement can be set at 1 nm (0.1 %) defocus spread, which corresponds to 0.23 V @ 200 kV and 0.35 V @ 300 kV. Such values will not be challenging to achieve electronically. The settling time of the modulation signal should be shorter than a single hardware frame of the detector. At present, the fastest direct detector on the market, the Gatan K3 (Gatan, Pleasanton, USA), operates at 1500 frames/s. Therefore, a settling time of <0.5 ms would be satisfactory and will not be difficult to achieve.

Future cryo-microscopes may be equipped with narrow gap objective lens polepieces, like the ones currently used in materials science. These lenses have ~3 times shorter focal length and similarly smaller chromatic aberration coefficient than current cryo-microscopes. Such systems will require proportionally higher biasing voltages on the aperture to produce equal defocus offsets. The objective lens aperture area may also be narrower, which will pose further practical difficulties. One possibility for implementing FADE on narrow gap lenses would be to voltage bias the sample instead of the aperture, but this would bring its own practical challenges. In any case, the motivation to develop and implement FADE on next generation systems will depend on the observed benefits, or lack thereof, after the method is tested in actual cryo-EM applications on current generation cryo-microscopes. We are working on performing such tests soon.

In this work, we focused our investigation on defocus modulation. Other optical parameters, such as beam tilt and astigmatism, should also be considered for real-time modulation. Such methods could offer additional optical and/or structural information. In general, we believe that the multiframe capability of direct detectors combined with real-time optical modulation has an untapped potential for further gains in cryo-EM performance.

Finally, the laser phase plate holds promise to become a permanent solution for the contrast vs resolution dilemma in cryo-EM. Until it has been perfected and becomes commercially available, or until the lack of resolution with strong defocus mystery has been resolved, the FADE method could provide an easy and cost-effective way to at least partially improve the performance.

## 5. Conclusions

In this study, we investigated the feasibility and performance of several types of modulators for the proposed FADE method. Electrostatic and magnetic micro-lenses were evaluated theoretically and numerically. The results showed that such devices will be difficult to construct to the required specifications. Using voltage bias on the objective lens aperture as a defocus modulator was also investigated. Results from simulations and experiments confirmed the feasibility and effectiveness of this approach. Aperture voltage modulation is easy to implement in practice and will become a straightforward and inexpensive way to expand the capabilities of existing cryo-microscopes. The FADE technique will offer an additional degree of freedom for tuning and optimization of cryo-EM experiments, for both single particle analysis and cryo-tomography. It has the potential to push the performance of cryo-EM a bit further along its development path.

## Acknowledgements

We are grateful to Masahide Kikkawa for helpful discussions and support with equipment and materials. We thank Aiko Shibata and Daiji Naka for their help with funding management. We are thankful to all members of the Quantum Biology JST PRESTO research area for stimulating discussions and helpful comments.

## Funding information

This work was supported by Japan Science and Technology Agency (JST) PRESTO grant #18069571.

